## supplemental figures 1-4 and supplemental tables for "Single-Cell Profiling Reveals Global Immune Responses during the Progression of Murine Epidermal Neoplasms"

Supplemental Figure 1

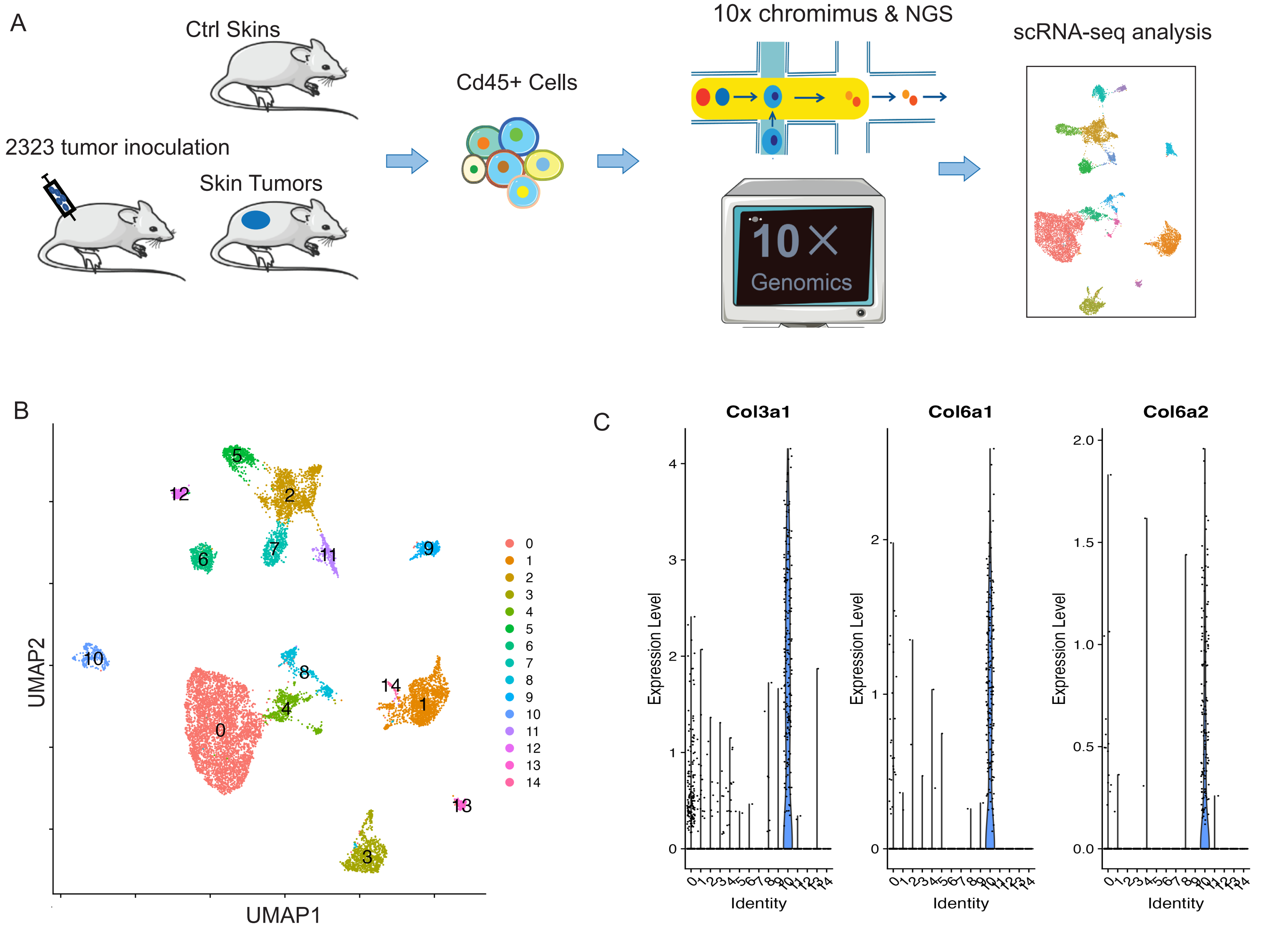

### Supplemental Figure 2

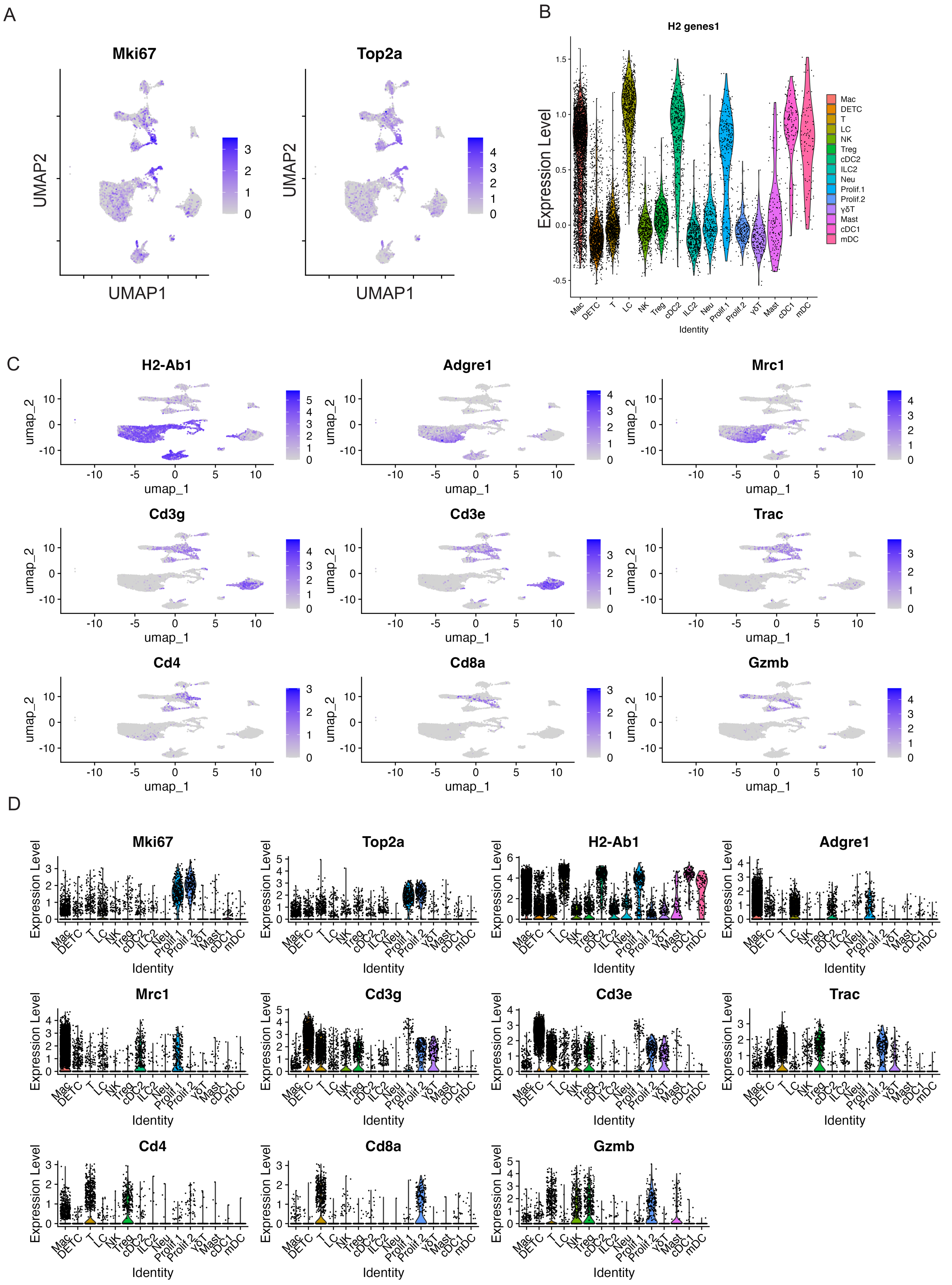

Supplemental Figure 3

A

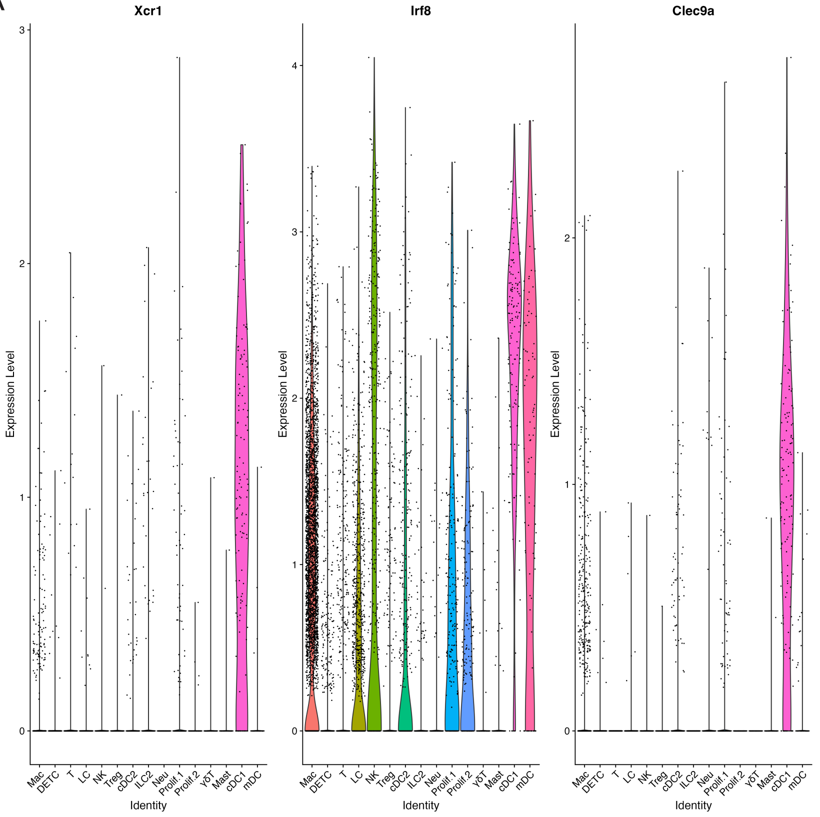

B

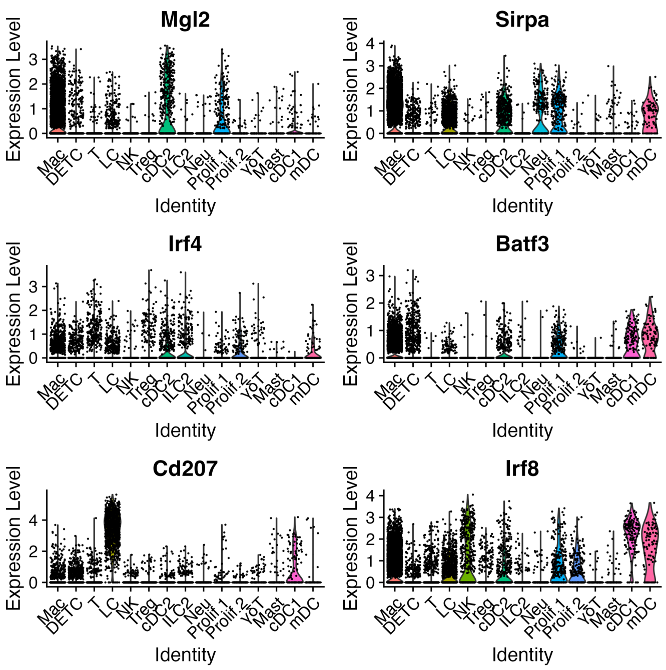

C

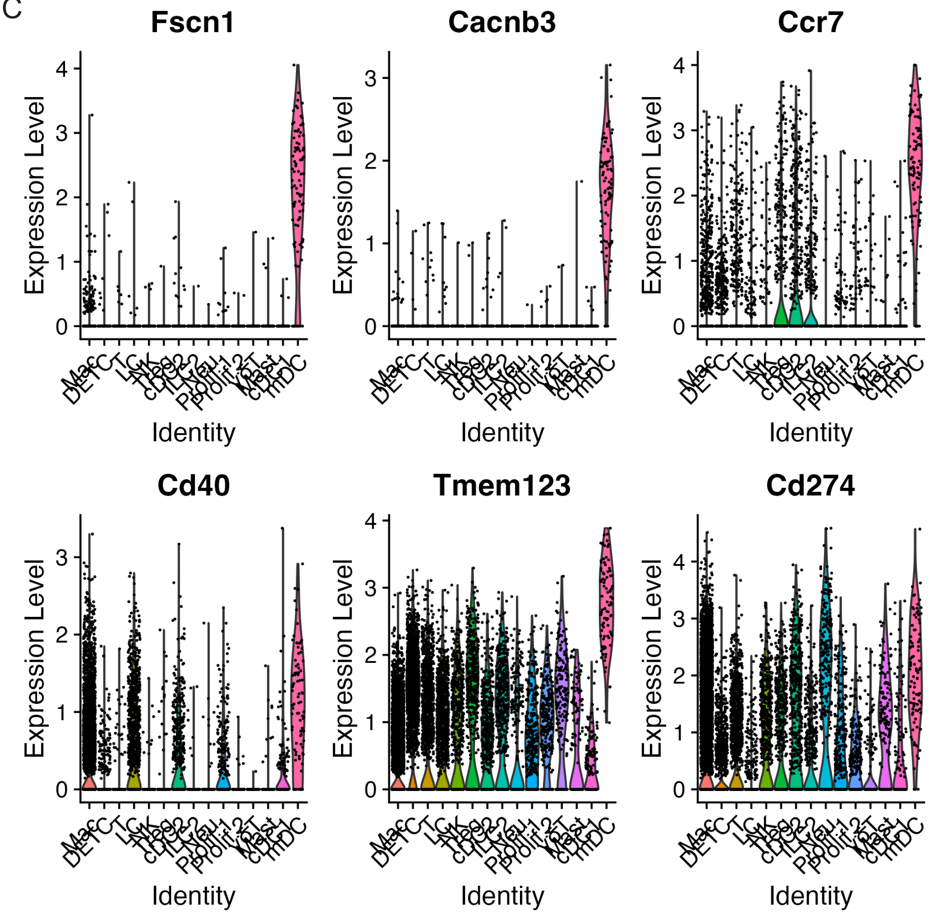

Supplemental Figure 4

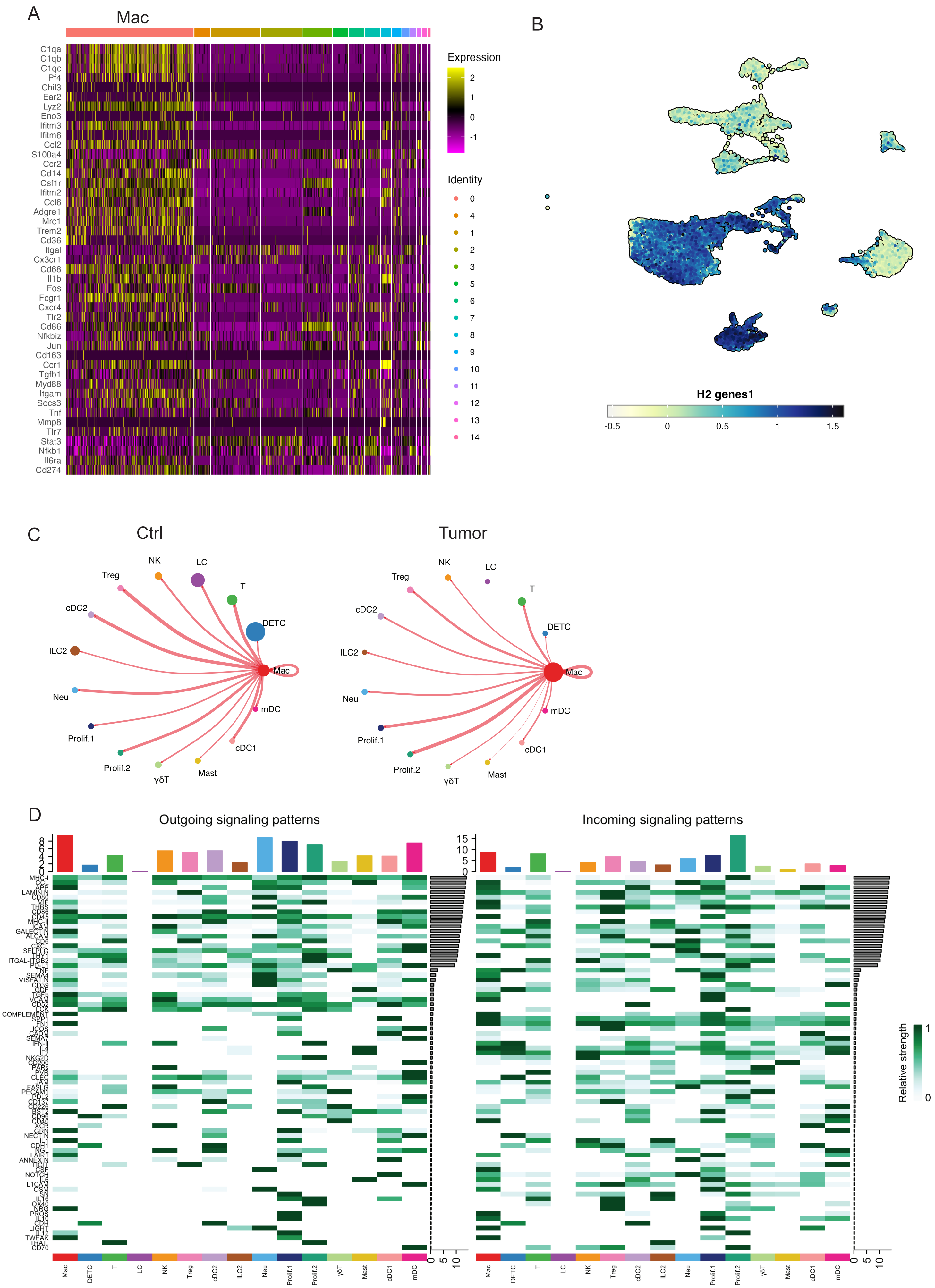

Table S1 scRNA-seq dataset metrics

| Samples | Estimated N | Post-filtering | Mean Reads per Cell |
| --- | --- | --- | --- |
| C57_1 | 3499 | 2898 | 87090 |
| C57_2 | 3862 | 3048 | 81098 |
| 2323_Tumor | 5176 | 4748 | 45977 |
| 2323_Tumor | 2258 | 1989 | 100120 |

Table S2 Representative marker genes for immune cell populations (related to Figure 1 and Figrue 2)

| | classic makers | % cell in cluster | Adjusted P-value | Extended markers | % cell in cluster | Adjusted P-value | Negative markers |
|---|---|---|---|---|---|---|---|
| Macrophage | Adgre1 | 69.6 | 0 | Mrc1 | 70.6 | 0 | |
| | Ms4a7 | 51 | 0 | Folr2 | 19 | 6.0111E-189 | |
| | C1qc | 69.5 | 0 | Arg1 | 12.6 | 2.1506E-148 | |
| | Itgam | 67.8 | 0 | | | | |
| | Cd68 | 77.8 | 0 | | | | |
| Type 2 conventional dendritic cells (cDC2) | H2-Ab1 | 98 | 2.13362E-85 | | | | |
| | H2-Aa | 96.2 | 2.60184E-78 | | | | |
| | Cd209a | 63.7 | 0 | | | | |
| | Cd209d | 30.6 | 7.3167E-224 | | | | |
| | Irf4 | 37.9 | 7.18645E-37 | | | | |
| | Mgl2 | 53.2 | 5.04279E-72 | | | | |
| Type 1 conventional dendritic cells (cDC1) | H2-Ab1 | 100 | 1.75795E-28 | | | | |
| | H2-Eb1 | 100 | 4.84001E-33 | | | | |
| | Clec9a | 82.8 | 0 | | | | |
| | Xcr1 | 78.6 | 0 | | | | |
| | Irf8 | 94.5 | 1.29475E-73 | | | | |
| | Batf3 | 67.6 | 5.66963E-43 | | | | |
| mature/migratory Dendritic cells (mDC) | Fscn1 | 87.1 | 0 | | | | |
| | Cancnb3 | 95.7 | 0 | | | | |
| | Ccr7 | 95.7 | 4.8695E-136 | | | | |
| | Cd40 | 82.8 | 5.2396E-34 | | | | |
| | Tmem123 | 100 | 3.29896E-53 | | | | |
| | Cd274 | 96.8 | 2.02954E-20 | | | | |
| Langerhans cells (LC) | H2-M2 | 95.4 | 0 | | | | |
| | Cd207 | 98.3 | 0 | | | | |
| | Epcam | 85 | 0 | | | | |
| | Cd24a | 71.5 | 0 | | | | |
| | Csf1 | 88.8 | 7.10E-214 | | | | |
| Dendritic Epidermal T cells (DETC) | Cd3e | 92.6 | 0 | | | | Cd4 |
| | Cd3g | 86.4 | 0 | | | | Cd8a |
| | Trdc | 95.5 | 0 | | | | |
| | Tcrg-C1 | 82.3 | 0 | | | | |
| | Thy1 | 76.9 | 0 | | | | |
| Dermal gamma delta T cells | Cd3e | 92.6 | 0 | | | | Cd4 |
| | Cd3g | 86.4 | 0 | | | | Cd8a |
| | Tcrg-C1 | 82.3 | 0 | | | | |
| | Trdc | 95.5 | 0 | | | | |
| | Thy1 | 76.9 | 0 | | | | |
| Natural Killer Cells (NK) | Gzma | 33.4 | 0 | | | | |
| | Klra7 | 40.5 | 0 | | | | |
| | Eomes | 42.6 | 0 | | | | |
| | Ncr1 | 40.3 | 0 | | | | |
| | Prf1 | 55.9 | 3.70E-140 | | | | |
| | Klra8 | 27.3 | 0 | | | | |
| | Klrb1c | 30.7 | 0 | | | | |
| Neutrophils (Neu) | S100a8 | 96.7 | 0 | | | | |
| | S100a9 | 97.6 | 0 | | | | |
| | Csf3r | 92.4 | 1.44E-297 | | | | |
| | Spi1 | 67.8 | 4.03E-18 | | | | |
| | Slc11a1 | 76 | 6.08E-117 | | | | |
| Mast Cells | Gata2 | 89.9 | 0 | Mctp4 | 98 | 0 | |
| | Ms4a2 | 62.4 | 0 | Kit | 88.6 | 2.93E-205 | |
| Type 2 innate lymphoid cells (ILC2) | Gata3 | 86.7 | 0 | | | Cd3d/e/g | |
| | Rora | 99.6 | 0 | | | Eomes | |
| | Il7r | 90 | 3.52E-207 | | | Rorc | |
| | Il5 | 48.1 | 0 | | | Tbx21 | |
| | Il13 | 41.1 | 5.79E-206 | | | Cd4 | |
| Treg | Foxp3 | 63.1 | 0 | | | | |
| | Cd4 | 42.8 | 0 | | | | |
| | Tnfrsf18 | 56.8 | 7.9504E-109 | | | | |
| | Ctla4 | 92.6 | 0 | | | | |
| Prolif.1 | Mki67 | 96.8 | 0 | | | | |
| | Top2a | 96.1 | 0 | | | | |
| | H2-Ab1 | 91 | 4.77347E-08 | | | | |
| | H2-Aa | 92 | 5.77427E-09 | | | | |
| | Adgre1 | 51.8 | 0.019728226 | | | | |
| | Mrc1 | 56.6 | 1.31607E-10 | | | | |
| Prolif.2 | Mki67 | 97.9 | 8.5769E-293 | | | | |
| | Top2a | 96.6 | 3.2286E-284 | | | | |
| | Cd3g | 82.9 | 2.48722E-51 | | | | |
| | Cd3e | 81.2 | 5.02767E-41 | | | | |
| | Trac | 80.8 | 1.0742E-140 | | | | |
| | Cd4 | 20.1 | 0.019835734 | | | | |
| | Cd8a | 58.1 | 5.3672E-225 | | | | |
| | Gzmb | 65 | 8.5022E-150 | | | | |
| T cells | Cd3d | 57.6 | 1.6044E-130 | | | | |
| | Trac | 62.8 | 0 | | | | |
| | Trbc1 | 45.4 | 2.15329E-61 | | | | |
| | Cd4 | 29.1 | 5.9231E-135 | | | | |
| | Cd8a | 33 | 0 | | | | |
| | Cd8b1 | 32.2 | 0 | | | | |

**Supplemental Figure 1. Our scRNA-seq experimental design**. (A) Graphical representation of the experimental setup. Samples were isolated from control skins and skin tumors. Live $CD45^+$ immune cells were FACS-sorted and loaded for scRNA-seq. (B) UMAP projections of 15 immune cell clusters at resolution 0.2. (C) Fibroblast genes Col3a1, Col6a1, Col6a2 genes highly expressed in cluster 10 indicating fibroblast contamination.

**Supplemental Figure 2. Two proliferating cell clusters**. (A) Feature plots showing two proliferative marker genes Ki67 and Top2a selectively expressed in two proliferating cell clusters. (B) Expression of MHC genes across immune cell clusters identifying APC cells such as Mac, LC, cDC1, cDC2, and mDC. (C) Feature plots showing the expression of H2-Ab1, Adgre1, Mrc1, Cd3g, Cd3e, Trac, Cd4, Cd8a, and Gzmb across immune cell clusters. (D) Violin plots of showing the expression of Mki67, Top2a, H2-Ab1, Adgre1, Mrc1, Cd3g, Cd3e, Trac, Cd4, Cd8a, and Gzmb across immune cell clusters.

**Supplemental Figure 3. DC marker genes across immune cell clusters**. The expression of cDC1 marker genes Xcr1, Irf8, and Clec9a across immune cell clusters. (B) Violin plots showing the expression of Mgl2, Sirpa, Irf4, Batf3, Cd207, and Irf8 across immune cell clusters. (C) Expression of mDC marker genes including Fscn1, Cacnb3, Ccr7, Cd40, Tmem123, and Cd274 across immune cell clusters.

**Supplemental Figure 4. Macrophage markers.** (A) A heatmap of macrophage markers showing the macrophage cell cluster. (B) Feature plot showing the H2 genes of APC clusters. (C) The circle diagram illustrates the signal crosstalk between macrophages and other immune cells, with the thickness representing the signal strength. (D) CellChat heatmap showing the relative strength of outgoing and incoming signaling pathways for each cell type. Vertical bars on top of the heatmap indicate overall importance of the cluster in the respective signaling pattern (i.e. outgoing or incoming). Horizonal bars indicate overall importance of the signaling pathway in the respective signaling pattern.
